# Quantitative mapping of membrane nanoenvironments through single-molecule imaging of solvatochromic probes

**DOI:** 10.1101/2020.07.19.209908

**Authors:** Daniel J. Nieves, Dylan M. Owen

## Abstract

Environmentally-sensitive fluorophores report on their local biochemical or biophysical environments through changes in their emission. We combine the solvatochromic probe di-4-ANEPPDHQ with multi-channel SMLM and quantitative analysis of the resulting marked point patterns to map biophysical environments in the mammalian cell membrane. We show that plasma membrane properties can be mapped with nanoscale resolution, and that partitioning between ordered and disordered regions is observed on length scales below 300 nm.

## Main Text

The mammalian cell plasma membrane is the site of processes essential for cellular function - homeostasis, mechanotransduction and signalling. Membrane biophysical properties play an integral role in regulating these processes. Once such property is membrane lipid order. It is hypothesised the plasma membrane can partition into disordered and ordered phase regions (Figure 1a)^1^. Ordered regions are enriched in saturated lipids and sterols, and are enriched in proteins which have high affinity for that environment. Ordered regions also display tighter lipid packing, thereby limiting the penetration of polar water molecules into the bilayer core. This feature can be exploited by solvatochromic probes that exhibit distinct changes in their fluorescence emission behaviour depending on the polarity of their surrounding environment^2^. Laurdan and di-4-ANEPPDHQ have previously been employed in this context using confocal microscopy, however, ordered domains are thought to exist below the resolution limit of conventional microscopy^3,4^.

**Figure 1:**
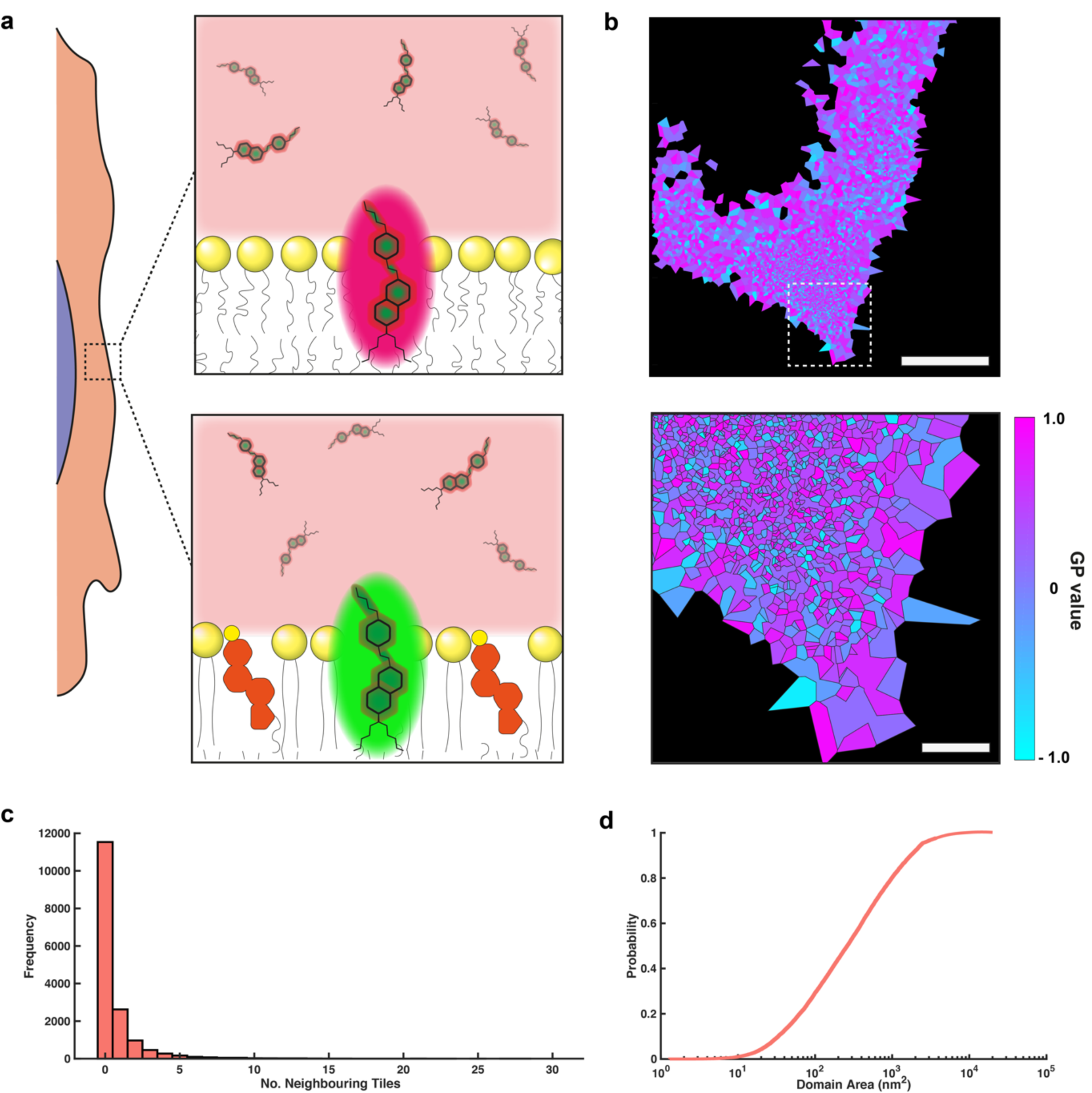
di-4-ANEPPDHQ-PAINT imaging of membrane domains. **a**) di-4-ANEPPDHQ in solution is free to bind to the cell membrane and, depending on the lipid environment, elicits a characteristic emission spectrum. **b**) Representative lipid order map generated via Voronoi tessellation and tile merging of a fixed RAMA27 fibroblast cell. Voronoi supertiles are colour-coded according to the average GP value for the tile. Scale bars 5 μm (top), 1 μm (bottom). **c**) Number of neighbouring tiles combined into each supertile. **d**) CDF of supertile areas.

Nile Red has previously been shown to be amenable to study by single-molecule localisation microscopy^5^. By recording the full emission spectrum for each localisation, changes in the emission wavelength due to differences in local environment can be observed^6,7^. Here, we demonstrate SMLM with the more commonly used dye, di-4-ANEPPDHQ^8^, to probe membrane lipid nano-environments in mammalian cell plasma membranes. di-4-ANEPPDHQ shows a large blue-shift in emission spectrum in ordered membrane regions when compared to disordered membrane regions (Figure 1a). Using a 2-channel imaging approach based on point accumulation in nanoscale topography (PAINT), we show that we can acquire the x-y coordinates of di-4-ANEPPDHQ insertion events into the cell plasma membrane. Furthermore, a ratiometric analysis of the two acquired spectral channels allows us to calculate a specific parameter for each point – the so-called Generalized Polarisation (GP) value – a measure of membrane order^8^; the resulting data, therefore, takes the form of a *marked point pattern*. We develop an analysis method for marked point pattern data, here in the context of membrane order, which has wide applicability for other marked data types.

di-4-ANEPPDHQ-PAINT was performed on fixed cells using 15 nM di-4-ANEPPDHQ in an oxygen scavenging buffer system (Glucose Oxidase:Glucose) to improve its photostability and making it amenable for SMLM imaging. Binding events at the plasma membrane were recorded simultaneously in the short and long wavelength channels (**Supplementary Movie 1**), and the sub-diffraction-limited positions of the individual binding events were extracted. Binding events were then matched frame by frame between the two channels, and the GP value calculated for each of the paired localizations using the extracted photon number (**Supplementary Figure 1**). The average position of the localisations in each channel is taken, and the GP value is ascribed to that position creating an *x, y*, GP coordinate. GP values run theoretically from -1 (highly disordered) to +1 (highly ordered). Using this approach, we were able to generate data with localisation precisions better than 30 nm (**Supplementary Figure 2**).

Voronoi tessellations have been implemented previously for both clustering and colocalization analysis of SMLM data. This turns a list of coordinates into a 2D set of tessellated polygons^9,10^, with the coordinates being at the centre of the polygons such that every position within one polygon is closer to that point than any other (**Supplementary Figure 3**). We implemented Voronoi tessellation on our GP SMLM data while retaining the GP value for each tile.

To visualise any ordered or disordered domains in the plasma membrane, we combined neighbouring polygons that had similar GP values (in this case, ± 0.1) into one supertile. We combined polygons until no more could be merged, thus giving a complete map of membrane domains within the data (**Figure 1b**). As a measure of domain size, we calculate the number of raw Voronoi polygons that are merged into each supertile (**Figure 1c**) and calculate the overall area of these tiles, displayed here as a cumulative probability distribution (**Figure 1d**).

While the Voronoi tessellations and generation of supertiles are useful for visualisation, they do not provide a statistical basis for the establishment of whether high or low order domains exist in the membrane. To achieve this, we applied a second approach, which interrogates the spatial separation, or colocalization, of the high and low order points. Given that we have points marked by a continuous range of GP values, (**Figure 2a**), we are able to split our localisations into “ordered” (**Figure 2b**, left) and “disordered” (**Figure 2b**, right) groups using a GP threshold. By splitting the data according to the membrane order, we can use analyses, which have been applied to conventional two-colour SMLM colocalization^11^, to determine whether high and low order points segregate from each other, or whether they are well-mixed. In this case, the GP threshold for splitting was set at 0.2.

**Figure 2:**
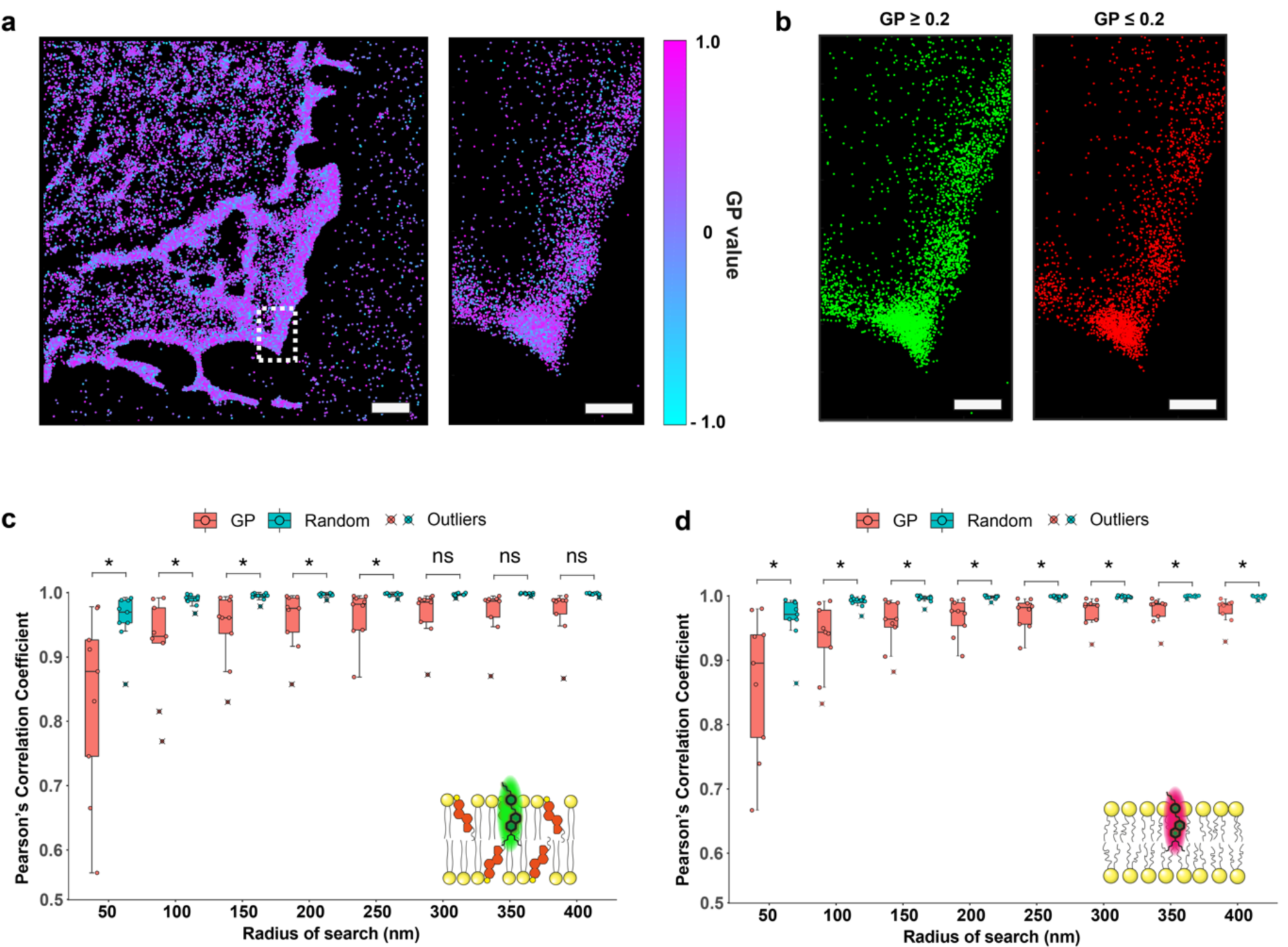
Lipid domains segregate at the nanoscale. **a**) Representative SMLM image of a RAMA27 fibroblast cell imaged using di-4-ANEPPDHQ-PAINT with points coloured by their GP value. Scale bar 2 μm (left), 500 nm (right). b) Points were segregated into two channels using the GP threshold of 0.2, corresponding to ordered (green) and disordered (red) channels. Scale bar 500 nm. **c-d**) *L*(r) vs *L*_*cross*_(r) analysis of the ordered and disordered channels respectively and compared to a random assignment of points into two channels. (ns: P > 0.05, **\***: P ≤ 0.05).

The localised versions of the Ripley’s K-function derivative, *L*(r), and the equivalent for the cross-K-function, *L*_*cross*_ (r), have been used previously to determine if point patterns in conventional two-channel SMLM data colocalise at a chosen length scale^11^ (**Supplementary Figure 4**). *L*(r) and *L*_*cross*_(r) were calculated for each point and plotted against each other, allowing the Pearson’s correlation coefficient to be calculated (**Supplementary Figure 4b and 4d**). We also performed the analysis on the same SMLM data, but this time randomly segregating points, irrespective of their GP value, into the two groups, as a control condition (**Supplementary Figure 4c and 4e)**. This process was then repeated for different values of the search radius, r, allowing the degree of colocalization to be quantified over different length scales.

For small search radii, the average calculated Pearson correlation coefficients were statistically significantly lower when segregating points by their GP values than those in the randomly assigned case (**Figure 2c and d**). However, for larger radii, there was no significant difference between the GP and random segregation showing that the low and high order points were well mixed at large radii. Thus, segregated ordered membrane regions exist only at a sub-diffraction spatial scale.

In summary, we have demonstrated that marked point pattern data can be extracted from the solvatochromic probe di-4-ANEPPDHQ using a 2-channel PAINT SMLM acquisition. When combined with statistical cluster and colocalization analysis, this can serve as a basis for probing and mapping the nanoscale spatial organisation of ordered lipid domains in the cell membrane. Our data suggest that large scale segregation of domains may not be commonplace, but that ordered domains can be observed segregated from disordered areas at the nanoscale (∼ 50-250 nm). This shows the potential for further membrane mapping approaches and lays the groundwork for analysis of other marked point patterns acquired by SMLM, for example using probes for membrane charge, pH, ion concentrations, viscosity or other parameters.

## Code Avaliability

Instructions and Matlab scripts to perform the analysis presented here are available at this repository: https://github.com/DJ-Nieves/di4ANNEPDHQ-PAINT.

## Author Contributions

DJN performed the experiments, designed and wrote the analysis codes, analysed data, and drafted and wrote the manuscript. DMO conceived the work, aided in analysis design, and wrote and drafted the manuscript.

## Acknowledgements

We acknowledge funding from BBSRC grant BB/R007365/1.

## SUPPLEMENTARY FIGURES

**Supplementary Figure 1:**
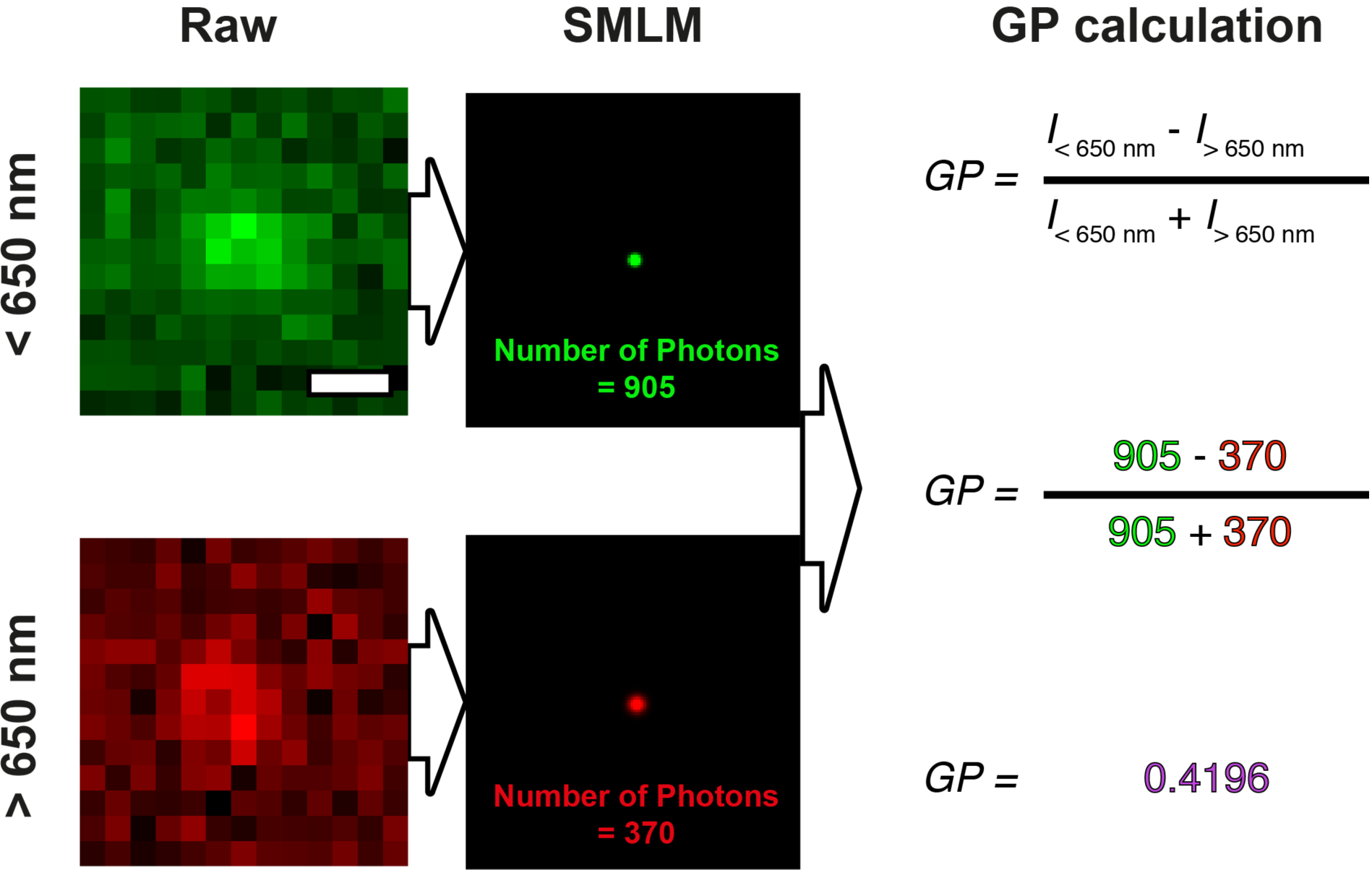
Processing and extraction of GP from SMLM data. di-4-ANEPPDHQ-PAINT binding events are recorded in long (> 650 nm) and short (< 650 nm) wavelength channels. The positions of the binding events and the number of photons per channel are determined by SMLM fitting. The photon numbers are then used to calculate the GP value for the event. Finally, the average position of the binding event is taken, and the GP value is ascribed to that point. Scale bar 500 nm.

**Supplementary Figure 2:**
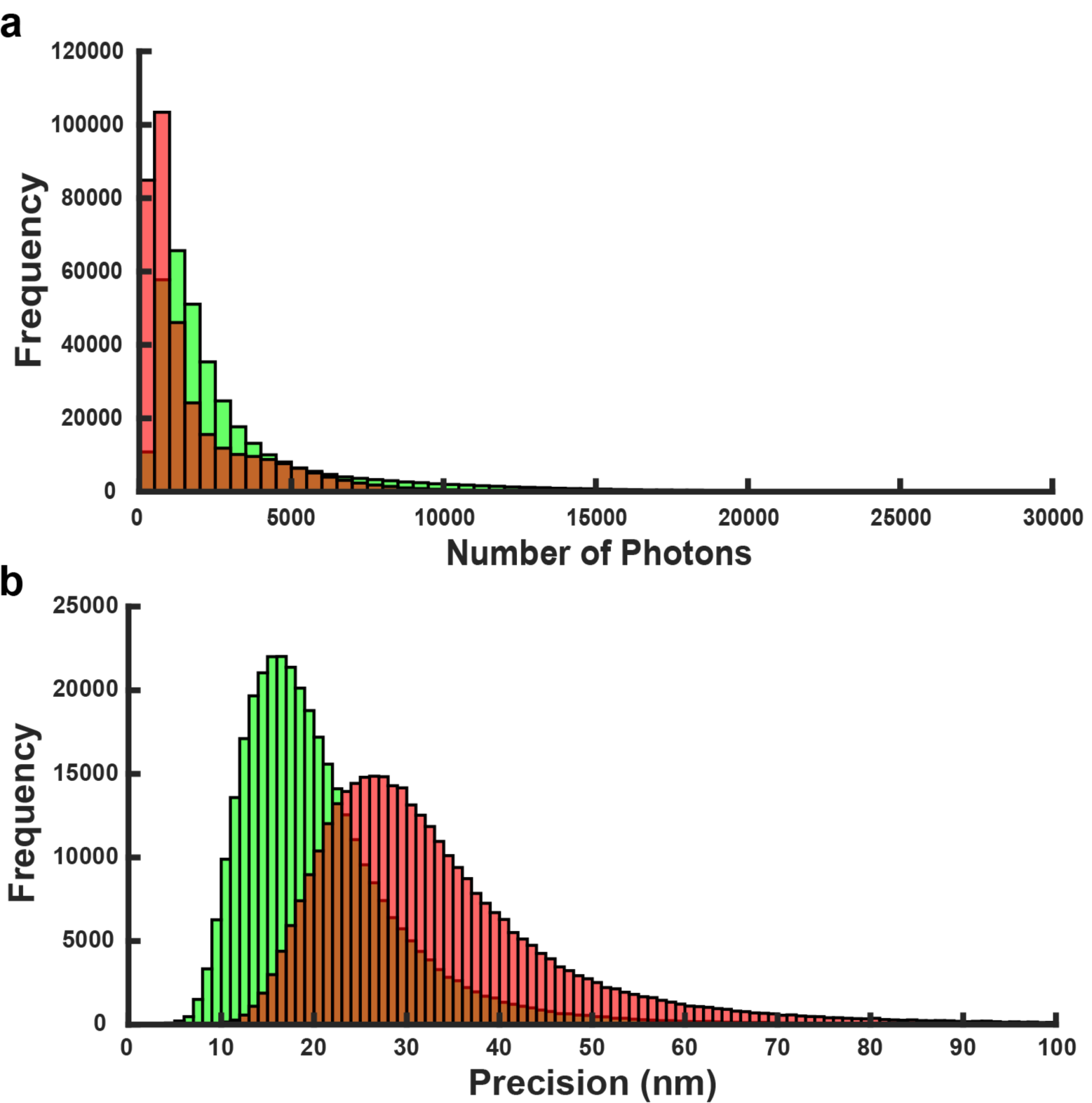
Photon count and precision histograms for paired GP SMLM data. **a)** Peak number of photons for paired binding events determined by SMLM fitting for each channel (< 650 nm: green, and > 650 nm: red). **b)** Calculated precision values for paired binding events for each channel (< 650 nm: green, and > 650 nm: red).

**Supplementary Figure 3:**
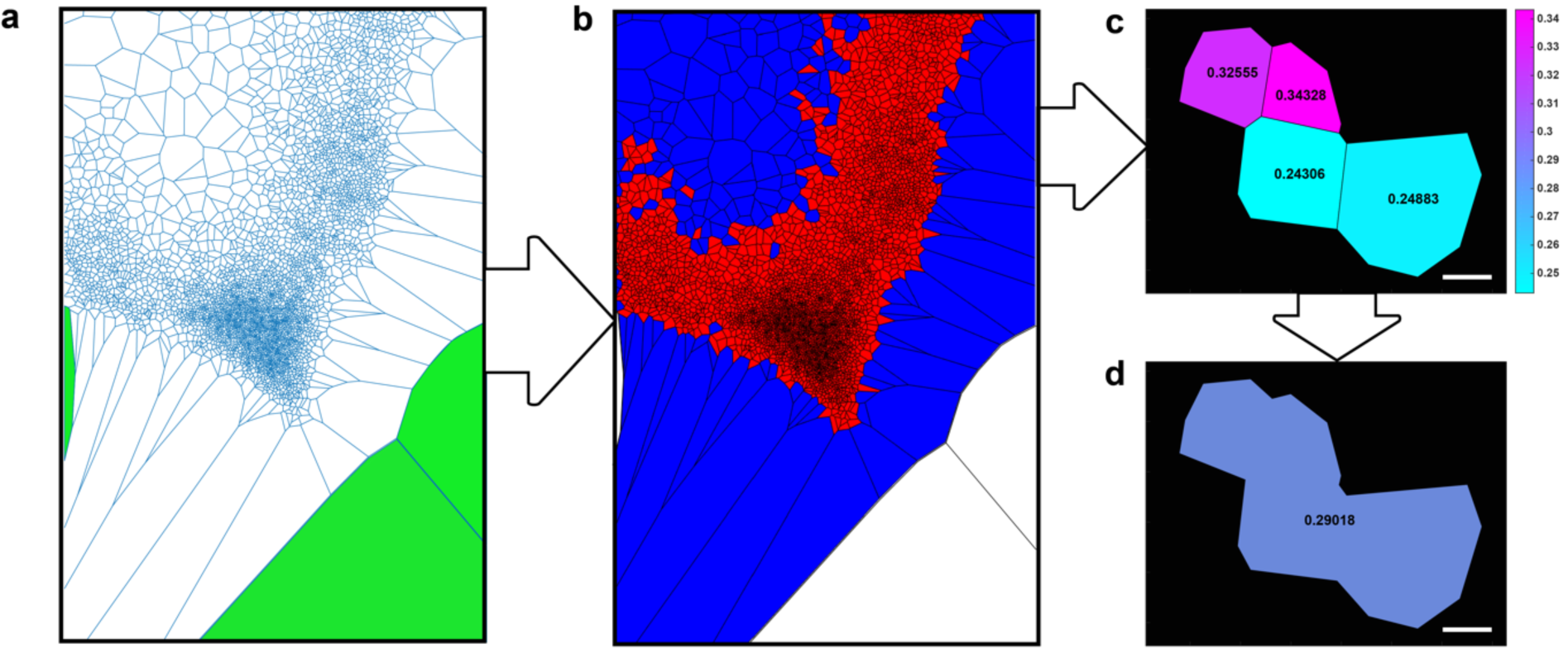
GP Voronoi Tessellation processing pipeline. **a**) The Voronoi diagram for the GP marked SMLM data is first computed, and tiles with infinity vertices, i.e., those at the edge of the region of interest, are discarded (green). **b**) The Voronoi tiles are then filtered to remove large tiles (greater than 2000 nm^2^), as these tiles represent areas of the data where di-4-ANEPPDHQ-PAINT event coverage and detection is low. **c**) Voronoi tiles are then labelled by their GP value. **d**) If neighbouring tiles GP values are within the GP tolerance range, they are concatenated into a larger tile, and the average GP value is calculated. Scale bar – 50 nm.

**Supplementary Figure 4:**
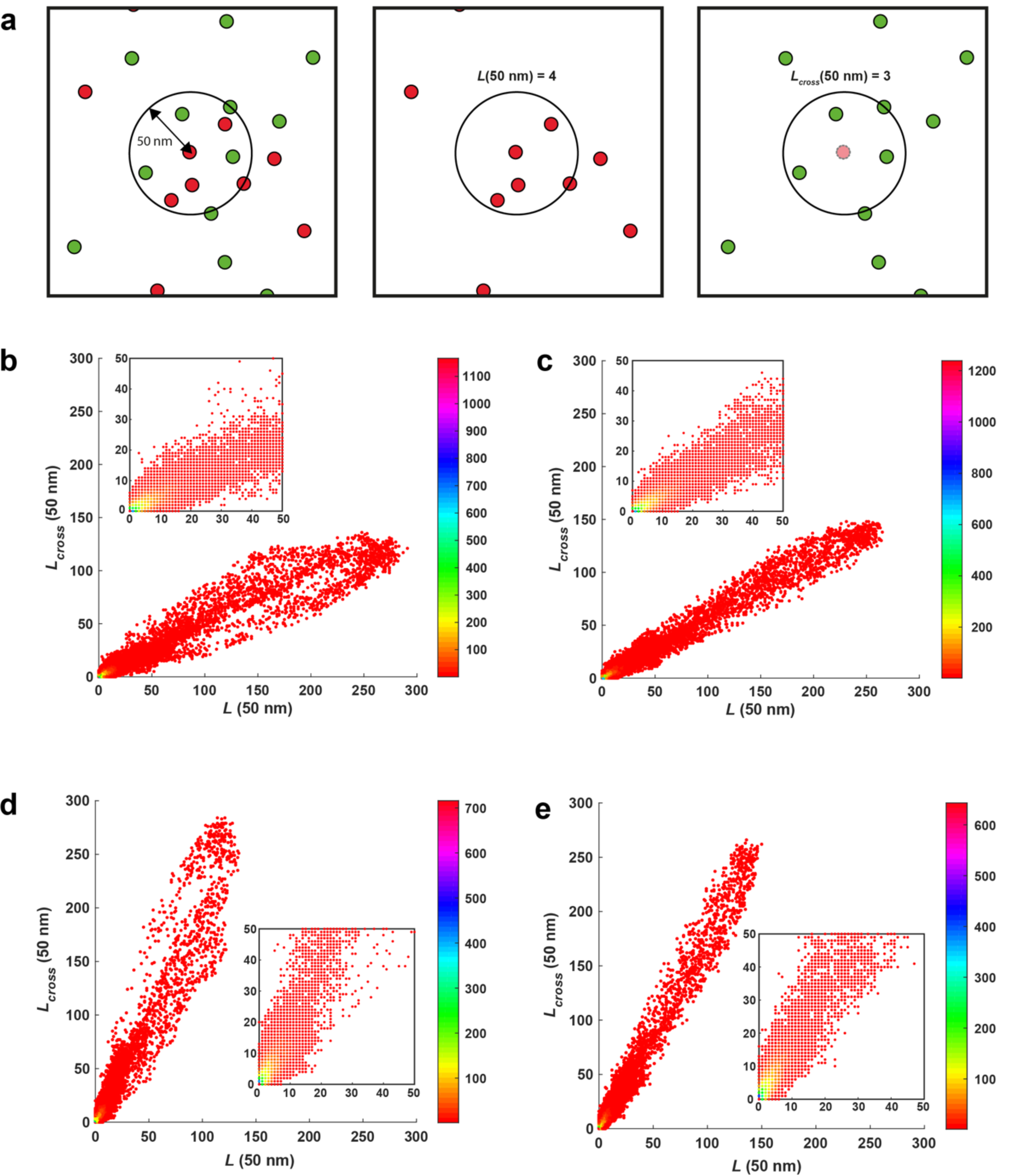
Principle and examples of colocalization analysis. **a**) Colocalization of two channel SMLM data can be determined by calculating the *L*(r) vs *L*_*cross*_(r) values for every point in a region of interest^11^. Two channel data is split, and the number of localisations within a radius of search (r, 50 nm in this example) is computed for every point, *L*(r) (middle). Next the number of localisations within the same radius is computed, but this time the search is for localisations in the opposing channel (right). **b**) Example *L*(r) vs *L*_*cross*_(r) for the GP > 0.2 channel. **c**) Example *L*(r) vs *L*_*cross*_(r) for randomly assigned data. **d**) Example *L*(r) vs *L*_*cross*_(r) for GP < 0.2 channel. **e**). Example *L*(r) vs *L*_*cross*_(r) for randomly assigned data. Colorbars represent the number of localisations possessing the specific *L*(r) vs *L*_*cross*_(r) value.

## SUPPLEMENTARY METHODS

### Cell culture and fixation

Rat mammary fibroblast cells (RAMA27), a gift from Prof. David G. Fernig (University of Liverpool), were cultured in plastic in Dulbecco’s modified Eagle’s medium (DMEM, Life Technologies, Paisley, UK) supplemented with 10% (v/v) fetal calf serum, 50 ng/mL insulin, 50 ng/mL hydrocortisone (all from Life Technologies) at 37°C in 10% (v/v) CO2 ^12^. When required for imaging they were trypsinized and seeded into Ibidi μ-slide 8-well glass bottomed chambers at 20,000 cells per well, and left over night in culture medium to adhere and spread. Cells were fixed prior to imaging with freshly prepared and warmed 4% PFA in PBS, from 16% methanol-free stock solution (Life Technologies).

### di-4-ANEPPDHQ-PAINT sample preparation

di-4-ANEPPDHQ (D36802; ThermoFisher Scientific) stock solution were prepared according to the manufacturer’s instructions, i.e., 1 mg/mL in DMSO and stored at 4°C. Prior to imaging a solution of 15 nM di-4-ANEPPDHQ was prepared in a glucose oxidase based scavenging buffer (10 mM Tris-HCl (pH 8), 50 mM NaCl, 0.8 mg/mL glucose oxidase, 18% (w/v) glucose). The 15 nM Di-4-ANEPPDHQ solution was added to the fixed cell samples just prior to imaging.

### di-4-ANEPPDHQ -PAINT imaging and image processing

Imaging was performed on an Oxford Nanoinstruments (ONI) Nanoimager. The sample was excited 488 nm, and the emission was split onto two halves the CMOS camera chip using a LP650 filter. Images were collected with an integration time of 50 ms for 20000 frames. Single molecule localisation of di-4-ANEPPDHQ binding events to the plasma membrane were fitted and extracted for each channel during the acquisition using the in-built ONI online fitting algorithms.

### di-4-ANEPPDHQ -PAINT data processing for GP analysis

Unfiltered single molecule localisation data from ONI online processing was first split into the short wavelength (< 650 nm) and long wavelength (> 650 nm) channels in MATLAB. The two channels were aligned post acquisition using image correlation by convolving the localisation data with a 2D gaussian and transforming the long channel image until high correlation spatial between the two channels was achieved. The transformation matrix from this process was then used to shift the localisation data in the long channel. Next, for each frame in the short channel the localisations were paired with corresponding localisations (within 100 nm radius) in the long channel. The photon numbers from each pair were taken, and used to calculate the GP value (**Supplementary Figure 1**);

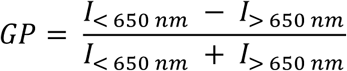

where *I*_< 650 *nm*_ is the photon count for the short wavelength channel and *I*_> 650 *nm*_ is the photon count for the long wavelength channel. This is process is repeated for all paired localisations. Finally, the average XY position of the localisations within the pair are taken to create a new coordinate, and the calculated GP value is ascribed to that point.

### GP Voronoi Analysis

The Voronoi diagram for the GP calculated localisation data was first calculated to yield tiles corresponding to each localisation. First, tiles possessing infinite coordinate values, i.e., those at the edges of the Voronoi diagram were removed. Then the tiles with areas smaller than 2500 nm^2^ were selected, as this represented the regions of the data with the best coverage from the di-4-ANEPPDHQ-PAINT. Next, for each tile, the number of neighbouring tiles (sharing more than one vertex) whose GP was within ± 0.1 of the GP of the tile of interest was calculated. If neighbours were found, the indexes of these tile were retained and grouped. The process of finding neighbours on the grouped tiles was repeated, and those groups, which shared neighbours were grouped. This was repeated until the number of tiles converged at a constant value, i.e., no new neighbouring’s could be found, thus confirming that complete tile grouping by GP value had been achieved. The combined area of these grouped tiles was then extracted and the mean GP value for the whole grouped tile domain calculated.

### *L*(r) vs *L*_*cross*_ (r) analysis

The GP calculated localisation data were split according to GP value, here, GP < 0.2 and GP > 0.2, to give two new localisations datasets. First the number of neighbouring points around each point within the same GP group were calculated within radii from 50 nm up to 400 nm to give the *L*(r). Then the search is repeated for each point, but this time counting the number of neighbours in the opposite GP channel giving the *L*_*cross*_ (r) for each point. This process was performed on 8 regions total from two cells. The *L*(r) vs *L*_*cross*_ (r) can then be plotted for each GP grouping (**Supplementary Figure 4 b-c**). The Pearson’s correlation coefficient was then calculated for *L*(r) vs *L*_*cross*_ (r) at every radius for each data region. This whole process was repeated for randomised data, i.e., localisation data from each region was split randomly into the two groups, rather than using the GP value (**Supplementary Figure 4 d-e**). The number of points in each group, however, was maintained for consistency with the GP sorting.

## References

1 Levental, I., Levental, K. R. & Heberle, F. A. Lipid Rafts: Controversies Resolved, Mysteries Remain. Trends Cell Biol 30, 341–353, doi: 10.1016/j.tcb.2020.01.009 (2020).

2 Klymchenko, A. S. Solvatochromic and Fluorogenic Dyes as Environment-Sensitive Probes: Design and Biological Applications. Acc Chem Res 50, 366–375, doi: 10.1021/acs.accounts.6b00517 (2017).

3 Pike, L. J. Rafts defined: a report on the Keystone Symposium on Lipid Rafts and Cell Function. J Lipid Res 47, 1597–1598, doi: 10.1194/jlr.E600002-JLR200 (2006).

4 Owen, D. M., Williamson, D. J., Magenau, A. & Gaus, K. Sub-resolution lipid domains exist in the plasma membrane and regulate protein diffusion and distribution. Nat Commun 3, 1256, doi: 10.1038/ncomms2273 (2012).

5 Sharonov, A. & Hochstrasser, R. M. Wide-field subdiffraction imaging by accumulated binding of diffusing probes. Proc Natl Acad Sci U S A 103, 18911–18916, doi: 10.1073/pnas.0609643104 (2006).

6 Danylchuk, D. I., Moon, S., Xu, K. & Klymchenko, A. S. Switchable Solvatochromic Probes for Live-Cell Super-resolution Imaging of Plasma Membrane Organization. Angew Chem Int Ed Engl 58, 14920–14924, doi: 10.1002/anie.201907690 (2019).

7 Bongiovanni, M. N. et al. Multi-dimensional super-resolution imaging enables surface hydrophobicity mapping. Nat Commun 7, 13544, doi: 10.1038/ncomms13544 (2016).

8 Owen, D. M., Rentero, C., Magenau, A., Abu-Siniyeh, A. & Gaus, K. Quantitative imaging of membrane lipid order in cells and organisms. Nat Protoc 7, 24–35, doi: 10.1038/nprot.2011.419 (2011).

9 Andronov, L., Orlov, I., Lutz, Y., Vonesch, J. L. & Klaholz, B. P. ClusterViSu, a method for clustering of protein complexes by Voronoi tessellation in super-resolution microscopy. Sci Rep 6, 24084, doi: 10.1038/srep24084 (2016).

10 Levet, F. et al. A tessellation-based colocalization analysis approach for single-molecule localization microscopy. Nat Commun 10, 2379, doi: 10.1038/s41467-019-10007-4 (2019).

11 Rossy, J., Cohen, E., Gaus, K. & Owen, D. M. Method for co-cluster analysis in multichannel single-molecule localisation data. Histochem Cell Biol 141, 605–612, doi: 10.1007/s00418-014-1208-z (2014).

12 Rudland, P. S., Twiston Davies, A. C. & Tsao, S. W. Rat mammary preadipocytes in culture produce a trophic agent for mammary epithelia-prostaglandin E2. J Cell Physiol 120, 364–376, doi: 10.1002/jcp.1041200315 (1984).

